# ACE 2 Coding Variants: A Potential X-linked Risk Factor for COVID-19 Disease

**DOI:** 10.1101/2020.04.05.026633

**Authors:** William T. Gibson, Daniel M Evans, Jianghong An, Steven JM Jones

## Abstract

Viral genetic variants are widely known to influence disease progression among infected humans. Given the recent and rapid emergence of pandemic SARS-CoV-2 infection, the cause of COVID-19 disease, viral protein variants have attracted research interest. However, little has yet been written about genetic risk factors among human hosts. Human genetic variation has proven to affect disease progression and outcome for important diseases such as HIV infection and malaria infestation. The fact that the human ACE2 protein is encoded on the X chromosome means that males who carry rare ACE2 coding variants will express those variants in all ACE2-expressing cells, whereas females will typically express those variants in a mosaic distribution determined by early X-inactivation events. This sex-based difference in ACE2 expression has unique implications for epidemiological studies designed to assess host genetic factors influencing progression from asymptomatic SARS-coV-2 infection to COVID-19. Here we present theoretical modelling of rare ACE2 coding variants documented to occur naturally in several human superpopulations and subpopulations, and show that rare variants predicted to affect the binding of ACE2 to the SARS-CoV-2 spike protein exist in people. Though the rs4646116 (p.Lys26Arg) allele is found in 1 in 70 Ashkenazi Jewish males, and in 1 in 172 non-Finnish European males, this allele is found at higher frequencies in females. Furthermore, the class of missense ACE2 alleles predicted to affect SARS-CoV-2 binding are found in aggregate among 1.43% and 2.16% of Ashkenazi males and females, respectively, as well as in 0.58% and 1.24% of European males and females outside of Finland. These alleles are rarer in other population groups, and almost absent from East Asians genotyped to date.

Though we are aware that full genome-wide and exome-wide sequencing studies may ultimately be required to assess human genetic susceptibility to SARS-CoV-2 fully, we argue on the basis of strong prior probabilities that genotyping of this class of alleles is justified in cases of atypical SARS-CoV-2 diseases, such as asymptomatic super-spreaders (if any are identified), and in neonatal/paediatric-onset COVID-19 disease. Even relatively rare susceptibility factors (1% or fewer carriers) may become quantitatively important in the context of hundreds of thousands of infections. A small number of asymptomatic carriers, or a small number of super-spreaders, or a small segment of the population that is disproportionately likely to require intensive care, can magnify the medical, social and economic impacts of a pandemic of this size. The speed of the pandemic and the large number of affected cases worldwide justify efforts to identify all possible risk factors for adverse outcomes, including efforts to identify genetic susceptibility factors in human hosts.

## Introduction

SARS-CoV-2 emerged as a human pathogen in late 2019.^1^ Though SARS-CoV-2^2^ infection is believed to be asymptomatic in some human hosts,^3^ COVID-19 (<u>co</u>rona<u>vi</u>rus <u>di</u>sease-20<u>19</u>)^4^ is a serious respiratory illness with fatality rates that vary from 1.4% of diagnosed cases^5^ to 50% or more among patients requiring ICU treatment.^6^ Age and preexisting comorbidities have been described as risk factors for more serious disease,^7, 8^ and environmental factors such as initial dose of virions and availability of critical care are also believed important. Early data from the pandemic has suggested that males might account for a greater proportion of cases^5^ and/or suffer from higher COVID-19 morbidity and mortality,^9^ though if this is ultimately shown to be true, the reasons behind it are unclear.

Given the recent and rapid emergence of the COVID-19 global pandemic, viral genetic and protein variants have attracted immediate early attention as putative risk factors for disease progression, though little has been written about genetic risk factors among human hosts. Human genetic variation has proven to affect disease progression and outcome for important infectious diseases; variation in CCR5 has been shown to affect the possibility of progression from HIV infection to AIDS,^10–14^ and human host genetic variation has long been recognized as mitigating the outcome of infestation by malarial parasites (reviewed by Kariuki and Williams^15^).

The human ACE2 protein has recently been shown to serve as the binding site and point-of-entry of SARS-CoV-2 into human cells.^16^ ACE2 had also previously been shown to bind SARS-CoV,^17^ a related virus that caused serious respiratory infections in humans in the early 2000s. The viral Spike glycoproteins (“S” proteins) of SARS-CoV^18^ and SARS-CoV-2^19^ bind physically to the peptidase domain of ACE2 using their receptor binding domains (RBD), thereby enabling viral internalization.

We note here that the *ACE2* gene is located on the X-chromosome; human males are hemizygous for the gene, expressing only one haplotype from their maternally-derived allele. Females, by contrast, have two X chromosomes and generally express both their paternal and maternal alleles in a mosaic distribution that is determined by early X-inactivation. Since we have prior knowledge that the ACE2 protein is bound directly by the viral S protein, and since prior protein-protein interaction modelling has identified specific amino acid residues within ACE2 that are bound directly by SARS-CoV-2, we propose here that coding variants at these specific sites are likely to be a genetic risk factor for progression from SARS-CoV-2 infection to COVID-19 disease, and we further propose that this risk would affect human males and females differently.

## Methods

The gnomAD database of human genetic variation catalogues coding variants from 141,456 adults without childhood-onset neurodevelopmental syndromes.^20^ Though little phenotypic detail is available on participants, this database serves as a convenient reference of “background” human genetic variation. We accessed the ACE2 entry in the gnomAD database (v.2.1.1) on March 30, 2020 and downloaded separate .csv files of the predicted missense variants and predicted Loss-of-Function (pLoF) variants in human ACE2. We manually counted the number of coding missense variants (242 variants) and pLoF variants (5 variants). We estimated sex-specific prevalences of rare and common coding ACE2 variants among the 76,702 male gnomAD participants (67,961 exomes and 8,741 genomes) by totalling hemizygotes for all variants, and among the 64,754 female gnomAD participants (57,787 exomes and 6,967 genomes) by totalling all variants, then subtracting the total number of hemizygotes and homozygotes for each variant, such that females homozygous for any variant were counted only once. We then obtained more granular allele count and frequency information from VCF files downloadable from gnomAD v2.1. We calculated Minor Allele Frequencies (MAFs) within each global superpopulation (e.g. Africans, South Asians) represented in gnomAD, and for specific subpopulations (e.g. Southern Europeans) where these were available. Among males, the prevalence of a rare X-linked variant is equivalent to the MAF for that allele. For females, the prevalence of a rare X-linked variant is equivalent to twice the MAF for that allele, because females have two X chromosomes (and hence, twice the number of X-linked alleles). We retrieved CADD scores from a downloaded version of CADD v1.4; the Phred-scaled CADD score^21^ is reported for position and allele combination. We also used the online tool LIST^22^ to estimate deleteriousness of coding amino acid substitutions in ACE2. Three-dimensional modelling used MolSoft.^23^ We also used the method of Schapira et al.^24^ to estimate the change in binding energy that would accrue for the 15 naturally-occurring missense variants.

## Results

### ACE2 is Generally Intolerant to Loss-of-function Variants

ACE2 is annotated as being intolerant to pLoF single-nucleotide variants (SNVs), with 3 pLoF SNVs observed, yet 31 such variants expected based on the size of the gene. The two stop-gain mutations (p.Leu116Ter and p.Leu656Ter) and the frameshift variant (p.Gly422ValfsTer15) are not seen among hemizygotes. The two splice variants that are seen in the hemizygous state may allow some correctly-spliced product to be made; allowing for that possibility, ACE2 variants confidently predicted to abrogate protein expression were not observed in the hemizygous state among the 76,702 male gnomAD participants (67,961 exomes and 8,741 genomes).

### ACE2 Missense Variants Occur at and Near Residues that Bind SARS-CoV-2

The gnomAD database catalogues 242 missense variants in ACE2, of which only 15 are predicted to lie at or near the Spike protein binding site of ACE2. The 67,961 male exomes and 8,741 male genomes in the gnomAD cohort sampled 76,702 X chromosomes, whereas the 57,787 female exomes and 6,967 female genomes sampled 64,754 X chromosomes (total number of X chromosomes originally sampled: 206,210). However, the actual sample size of chromosomes for each variant is smaller, because not all X chromosomes genotyped successfully at each site (Supplementary Table 1). The range varied from 158,104 X chromosomes genotyped successfully for p.Val488Ala to 183,374 X chromosomes that genotyped successfully for p.Glu35Lys. Variant frequencies in Supplementary Table 1 were calculated using the number of alternate allele counts as the numerator and the number of chromosomes genotyped successfully at the locus as the denominator. Aggregating the allele frequencies of ACE2 variants located at the SARS-CoV-2 binding site (i.e. excluding p.Asn720Asp because it is not at the binding site) yields a frequency of 0.0041, which estimates that roughly 3.9 males per 1000 and 8.5 females per 1000 would harbour a rare allele that might affect the binding of SARS-CoV-2 to cells that express ACE2. Importantly, these rare alleles are not distributed evenly across human subpopulations, as outlined below.

The only common missense Single-Nucleotide Polymorphism (SNP) is rs41303171, predicted to encode p.Asn720Asp (UniProt Accession Number Q9BYF1). This SNP was observed in 1.7% of male gnomAD samples and in 3.1%-3.2% of female participants overall. Calculated allele frequencies among females vary slightly depending on how the counts are done. The downloadable .csv file lists the rs41303171 variant allele as observed 3,054 times: from that we can subtract 1,026 hemizygotes and 34 homozygotes (to avoid counting those females twice), yielding 1,994 heterozygotes + homozygotes among 64,754 female samples, such that 3.08% of female samples are estimated to have at least one p.Asn720Asp. This method assumes that all samples genotyped successfully at that site. To take genotyping efficiency into account we downloaded the relevant .vcf files and divided the Alternate Allele Count (1,809) by the Allele Number at that locus (112,288 alleles genotyped successfully among females) to yield a MAF of 1.6%, so 3.2% of females have at least one p.Asn720Asp allele. This allele would be frequent enough to be detected by Genome-Wide Association Studies (GWAS), either directly or by imputation, if this variant or one nearby were involved in the disease under study; to date no GWAS have flagged this locus.^25^ However, rs41303171 does not lie in a domain of the protein that is predicted to interact with SARS-CoV-2, so it is not an obvious candidate site for this particular host-pathogen interaction.

The various superpopulations and subpopulations in gnomAD each offer different sample sizes. For rare X-linked variants it is best to deconvolute allele frequencies by both sex and subpopulation, to achieve a fuller picture of the prevalence of hemizygous males in individual regions or countries. Any disease risk signal conferred by a functional X-linked allele will otherwise be diluted by lumping hemizygous males in with heterozygous female carriers, who are likely at a lower risk due to mosaic expression of the variant. After rs41303171, rs4646116 is the next-most-common missense ACE2 variant (global MAF 0.4%); it is predicted to encode p.Lys26Arg. Our 3D modelling predicted p.Lys26Arg to lie immediately beside the ACE2-Spike protein interface (albeit with the lysine side chain pointing away from the interface). Prior work on ACE2 binding to SARS-CoV^18, 26^ and recent work on ACE2 binding to SARS-CoV-2^27, 28^ by others has identified three main S protein binding domains within ACE2. The largest includes Gln24, Thr27, Phe28, Asp30, Lys31, His34, Glu35, Asp38, Tyr41, Gln42 and Leu45. The smallest comprises Met82 and Tyr83, and the last includes Asn330, Lys353, Gly354, Asp355 and Arg357. According to Li et al,^29^ residues Asp30 to Tyr41 make up the alpha-1 ridge, residues Met82 to Val93 make up the Loop and alpha-3 ridge, and residues Lys353 to Arg357 make up the Loop and beta-5-helix.

We estimated a high prior probability that human germline coding variants affecting these key binding residues would also affect the affinity of the endogenous human ACE2 protein for the viral spike protein. By visual inspection of gnomAD’s ACE2 missense variant list, we conservatively considered p.Lys26Glu, p.Lys26Arg, p.Thr27Ala, p.Glu35Lys, p.Glu37Lys, p.Phe40Leu, p.Ser43Arg, p.Met82Ile, p.Pro84Thr, p.Gly326Glu, p.Glu329Gly, p.Gly352Val, p.Asp355Asn and p.Val488Ala to be at or near the ACE2-S protein interface. All of these variants occur naturally in humans, and could plausibly affect risk for progression to COVID-19 after an initial encounter with SARS-CoV-2. We modelled the interface between ACE2 and the viral spike protein as shown in Figure 1.

**Figure 1:**
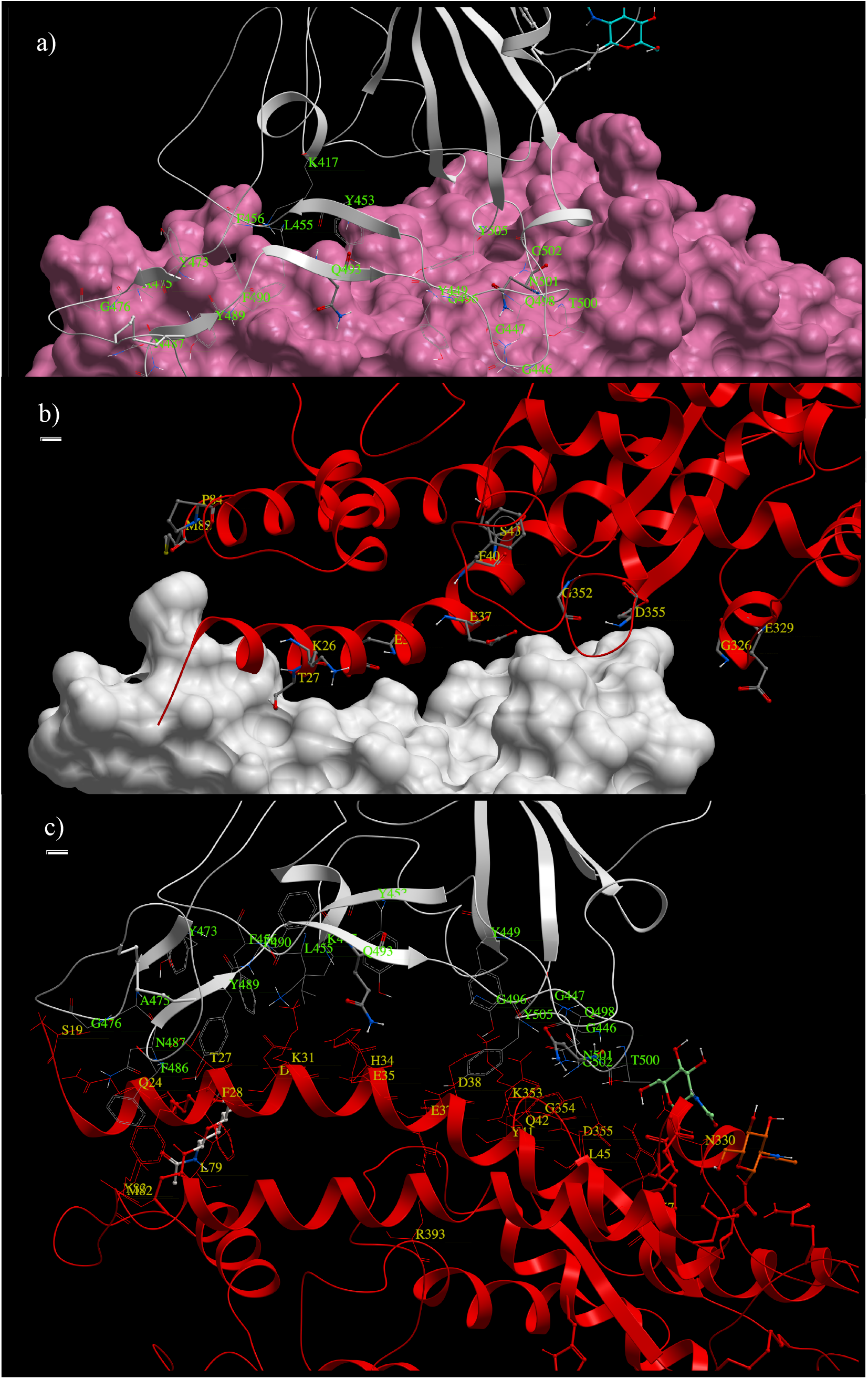
Modelling of the interface between human ACE2 and SARS-CoV-2 spike protein. The SARS-CoV-2 spike protein is presented with amino acid residues labelled in green, and ACE2 residues labelled in yellow. The spike protein is presented in white (ribbon or density map). ACE2 is presented as a pink density map (a) or red ribbon (b,c). The ACE2 residues that interact directly with viral Gln493 are predicted to be Lys31, His34 and Glu35. The ACE2 residues that interact directly with viral Asn501 are predicted to be Tyr41, Lys353 and Asp355. Apart from Asp355 and Glu35, additional ACE2 residues that are predicted to bind directly to the SARS-CoV-2 spike protein are Thr27, Glu35, Glu37, Met82 and Gly326.

Considering this class of ACE2 variants as a group, their estimated aggregate prevalence is 3.9 per 1000 males and 8.5 per 1000 females. Rare deleterious X-linked alleles will be depleted in live male adults, and observed disproportionately among heterozygous carrier females. Thus, the observation that this class of alleles is more than twice as common among females as in males may reflect some degree of historical selection against these alleles. We went on to calculate subpopulation-specific frequencies by sex, as .vcf files downloadable from the gnomAD interface provide the breakdown of exomes and genomes by subpopulation and by sex. rs4646116 (p.Lys26Arg) is found at polymorphic frequencies (MAF 1.43%, or 1 in 70) in Ashkenazi Jewish males, and at lower frequency among non-Finnish European males (MAF 0.58%, or 1 in 172). In Southern Europeans, not segregated by sex, the MAF is 0.34% (1 in 300). In males of Latino ancestry the MAF is 0.28% (1 in 360), and the frequency is lower among African Males (0.13% or 1 in 770), South Asian males (0.08% or 1 in 1,250) and Finnish males (0.07% or 1 in 1,430). This allele is at very low frequencies in East Asian participants, being observed only once in a heterozygous female (MAF of 0.01%). The allele was absent from Korean (n=2887 chromosomes) and Japanese (n=122 chromosomes). The remaining alleles p.Lys26Glu, p.Thr27Ala, p.Glu35Lys, p.Glu37Lys, p.Phe40Leu, p.Ser43Arg, p.Met82Ile, p.Pro84Thr, p.Gly326Glu, p.Glu329Gly, p.Gly352Val, p.Asp355Asn and p.Val488Ala are encountered very infrequently (between 0 and 4 times in each of the gnomAD subpopulations). However, since all of these naturally-occurring alleles occur at the binding interface, they may legitimately be grouped together to estimate allelic burden arising from this class of alleles. The fact that rare X-linked alleles are observed twice as often in females means that 2.1% of Ashkenazi Jewish females (1 in 48), carries a missense variant predicted to affect binding of SARS-CoV-2, as do 1.2% of non-Finnish European females. Similar calculations show that 0.74% of Latino females (1 in 135), 0.48% of South Asian females (1 in 208), 0.28% of African females (1 in 360) and 0.13% of Finnish females (1 in 770) carry a rare ACE2 allele that may affect binding to SARS-CoV-2.

The CADD scores among this class of alleles range from 0.017 to 24.9. We provide these scores here for interest. LIST scores for predicting the deleteriousness of all possible missense ACE2 alleles are available at https://list.msl.ubc.ca/proteins/Q9BYF1. We offer the caveat that scores of deleteriousness are difficult to interpret in this context, because they are not based on specific characteristics of a protein’s ability to bind to coevolved (or in this case, newly-evolved) partners.

### Human ACE2 Missense Variants May Increase or Decrease its Binding to SARS-CoV-2

Applying the method of Schapira et al.,^24^ we estimated the change in binding energy of the 15 missense variants (Supplementary Table 2). Glu37Lys increased the binding the most, followed by Thr27Ala, Lys329Gly, and Lys26Glu. Val488 and Asn720 are relatively distant from the interface, so they may contribute to the binding of other partners, but not likely the S protein. Several variants were predicted to weaken binding between ACE2 and SARS-CoV-2 spike protein: Asn720Asp weakened binding somewhat, with Ser43Arg, Gly326Glu, Met82Ile and Lys26Arg all weakening binding progressively more. The fact that Lys26Glu and Lys26Arg seem to have opposing effects suggests that the bulkier arginine side chain may not pack as tightly with viral S protein, whereas replacing the basic lysine with an acidic glutamate residue may disrupt important polar interactions.^30^ Brielle et al. note that the Phe486 of SARS-CoV-2 is critical for binding ACE2 residues Leu79 and Met82, thereby stabilizing the S protein-ACE2 interaction.^31^ Methionine 82 interacts with Phe486 of the viral protein’s RBD via van der Waals forces, so an isoleucine at this position may weaken this interaction.^30^ Yan et al. note that Arg426 of SARS-CoV forms a salt bridge with Glu329 on ACE2, and that SARS-CoV-2 has Asn439 at the analogous position, which cannot form this salt bridge. We speculate here that replacing glutamate 329 with a smaller glycine side chain in the p.Glu329Gly variant may allow easier binding of the SARS-CoV-2 spike protein’s Asn439, though this remains to be tested.

## Discussion

At the time of this writing, SARS-CoV-2 has infected over 1,250,000 people, with over 335,000 cases in the United States, approximately 130,000 cases in each of Italy and Spain, over 100,000 cases in Germany and nearly as many again in France. China has reported 3,333 deaths among 82,602 cases;^32^ since new cases have dropped precipitously there, we may estimate a fatality rate among recognized cases at 4%. However, Spain has reported over 12,500 deaths, and Italy has reported nearly 16,000 deaths. Though a large number of asymptomatic and/or unreported cases would dilute the true case-fatality rate, fatality rates of 9.6% and 12% among the subset of cases that are recognized and reported are significant. The speed of the pandemic and the large number of affected cases worldwide justify efforts to identify all possible risk factors for adverse outcomes, including efforts to identify genetic susceptibility factors in human hosts. Even relatively rare susceptibility factors (1% or fewer carriers) may become quantitatively important in the context of hundreds of thousands of infections. A small number of asymptomatic carriers, or a small number of super-spreaders, or a small segment of the population that is disproportionately likely to require intensive care, can magnify the medical, social and economic impacts of a pandemic of this size.

The genetic component of host-pathogen interaction studies forms a subset of gene-by-environment interaction studies. These latter studies require 10^5^ or more participants and sensitive measures of environmental exposures, and typically identify tagging SNPs linked to loci with small effect sizes. These methods, their attendant complexities, and the conservative genomewide thresholds they require are justified by the fact that previous candidate-gene studies had extremely high false-discovery rates,^33^ largely due to the use of convenience samples that generated artefactual signals owing to population stratification (i.e. the fact that SNP allele frequencies vary between ancestral populations for historical reasons that are unrelated to heritable risk for disease). However, given the truly staggering number of SARS-CoV-2 infections and the rapid progression from COVID-19 to death (often within 14 days), such labour-intensive gene-agnostic methods are unlikely to yield clinically-actionable results during the current pandemic.

For these reasons, we argue that prior knowledge of SARS-CoV biology and emerging knowledge of SARS-CoV-2 similarities to SARS-CoV justify the assessment of variation within human ACE2 as an exceptionally strong candidate gene for host response to SARS-CoV-2 infection. The fact that ACE2 is X-linked means that rare variants that enhanced SARS-CoV-2 binding *in vivo* would likely increase susceptibility to COVID-19 among males. Other functional effects are possible also; for example, female variant carriers might have a shorter asymptomatic period, or be more likely to develop symptoms, irrespective of viral shedding status. Given that a limited number of coding variants in ACE2 are predicted to affect the binding of SARS-CoV-2 spike protein, these variants could be genotyped rapidly in early-onset cases (e.g. paediatric cases, or even cases under age 40) as a pilot study in the United States and/or in European countries where case numbers are high, case fatality appears to be high and the rare alleles are likely to be found. The delivery of viral nucleic acid testing to affected populations is a more urgent need, but given that many viral samples are already being collected, the addition of a rapid assay targeting the SNVs listed here would be an efficient means to gather data to test this hypothesis. Notably, tests of rare variant burden do lose power rapidly if neutral variants and variants with opposing functional effects are grouped together, so we have restricted ourselves to variants with prior evidence of involvement in SARS-CoV-2 Spike protein binding. There are many more missense ACE2 variants in gnomAD; the aggregate crude prevalence of all rare (i.e. excluding the common p.Asn720Asp) missense variants in gnomAD males is 2.49% and in gnomAD females is 5.81%. Furthermore, among patients with extreme outcomes such as death from COVID-19 in childhood (or asymptomatic super-spreaders, if any are found), sequencing of the entire ACE2 coding region might conceivably identify ultra-rare “private” variants not catalogued in gnomAD, enabling assessment of the role of these variants in the conversion from initial SARS-CoV-2 infection to COVID-19. A properly-powered genome-wide study with correlation of genetic variants to multiple other clinical risk factors and outcomes is also advisable, but cannot realistically be accomplished in the near term.

Procko has done unbiased mutagenesis of the ACE2 regions that bind to the RBD of SRS-COV-2,^34^ and noted that substitution of Thr27 with hydrophobic residues (e.g. alanine) is expected to increase the ability of aromatic residues of the S protein to pack into the ACE2 interface. Procko’s *in vitro* experiments flagged Asn 90 and Thr92 as sites at which nonsynonymous variants should increase binding of the viral RBD motif. The p.Thr92Ile variant (rs763395248) occurs naturally in Southern Europeans (2 out of 7040 X chromosomes in gnomAD, both in males, MAF of 2.8 x 10^−4^). This *in vitro* work offers proof-of-principle for our hypothesis that naturally-occurring human ACE2 variants are likely to affect SARS-CoV-2 infection kinetics *in vivo*. At the very least, further *in vitro* functional assays of viral binding and replication within cell lines transfected to express human variant ACE2 proteins are urgently justified. Furthermore, given recent data that human recombinant soluble ACE2 (hrsACE2) can block early stages of SARS-CoV-2 infections,^35^ any naturally-occurring human ACE2 variant that bound more tightly to the viral spike protein might serve as a better “decoy” for the viral protein, and would be a candidate for novel hrsACE2 molecules with therapeutic potential.

Whether or not these studies can be done in time to affect patient outcomes during the current pandemic, the fact that three pathogenic coronaviruses (SARS-CoV,^36^ MERS-CoV^37^ and SARS-CoV-2^38^) have emerged in the past two decades, two of which bind and enter via ACE2, augurs that additional coronaviruses that bind to ACE2 are likely to emerge in the future. Thus, knowledge of this specific host-pathogen interaction at the molecular level is important to have at this time.

