## Supplementary Table 1 for "ACE 2 Coding Variants: A Potential X-linked Risk Factor for COVID-19 Disease"

| X Chromosome Position (GRCh37, hg19 build) | rsID | Reference Allele | Alternate Allele | Predited Protein Consequence | Transcript Consequence | CADD Score | Alternate Allele Count (AC) | Total Allele Number (AN) | Minor Allele Frequency (MAF) (among all chromosomes successfully genotyped at that position) | Allele Count in the population with the highest MAF | Total number of alleles in the population with the maximum MAF | Maximum observed MAF across all genotyped populations (excluding samples of Ashkenazi, Finnish, and unknown ancestry) | Alternate Allele Count (All Males in gnomAD Database) | Total Allele (All Males in gnomAD Database) |
|---|---|---|---|---|---|---|---|---|---|---|---|---|---|---|
| 15618959 | rs1299103394 | T | C | p.Lys26Glu | c.76A>G | 11.44 | 1 | 183,326 | 5.45E-06 | 1 | 81,870 | 0.0012% | 1 | 67,760 |
| 15618958 | rs4646116 | T | C | p.Lys26Arg | c.77A>G | 10.49 | 728 | 183,330 | 3.97E-03 | 483 | 81,876 | 0.5899% | 259 | 67,766 |
| 15618956 | rs781255386 | T | C | p.Thr27Ala | c.79A>G | 12.53 | 2 | 183,319 | 1.09E-05 | 2 | 27,385 | 0.0073% | 0 | 67,753 |
| 15618932 | rs1348114695 | C | T | p.Glu35Lys | c.103G>A | 4.551 | 3 | 183,374 | 1.64E-05 | 2 | 13,847 | 0.0144% | 1 | 67,810 |
| 15618926 | rs146676783 | C | T | p.Glu37Lys | c.109G>A | 23.7 | 6 | 183,339 | 3.27E-05 | 1 | 13,163 | 0.0076% | 3 | 67,777 |
| 15618915 | rs924799658 | G | T | p.Phe40Leu | c.120C>A | 6.419 | 3 | 183,233 | 1.64E-05 | 3 | 27,349 | 0.0110% | 0 | 67,671 |
| 15618906 | rs1447927937 | A | T | p.Ser43Arg | c.129T>A | 22 | 1 | 183,088 | 5.46E-06 | 1 | 18,988 | 0.0053% | 0 | 67,526 |
| 15613067 | rs766996587 | C | T | p.Met82Ile | c.246G>A | 0.017 | 2 | 182,758 | 1.09E-05 | 2 | 13,148 | 0.0152% | 0 | 67,236 |
| 15613063 | rs759134032 | G | T | p.Pro84Thr | c.250C>A | 3.543 | 1 | 182,792 | 5.47E-06 | 1 | 18,823 | 0.0053% | 0 | 67,270 |
| 15613038 | rs763395248 | G | A | p.Thr92Ile | c.275C>T | 2.403 | 2 | 182,553 | 1.10E-05 | 2 | 7,040 | 0.0284% | 2 | 67,019 |
| 15599437 | rs759579097 | C | T | p.Gly326Glu | c.977G>A | 12.88 | 1 | 181,266 | 5.52E-06 | 1 | 13,058 | 0.0077% | 0 | 65,872 |
| 15599428 | rs143936283 | T | C | p.Glu329Gly | c.986A>G | 12.24 | 5 | 181,401 | 2.76E-05 | 4 | 81,369 | 0.0049% | 0 | 65,981 |
| 15599359 | rs370610075 | C | A | p.Gly352Val | c.1055G>T | 24.9 | 1 | 173,847 | 5.75E-06 | 1 | 78,468 | 0.0013% | 0 | 59,229 |
| 15599351 | rs961360700 | C | T | p.Asp355Asn | c.1063G>A | 23.8 | 2 | 170,310 | 1.17E-05 | 2 | 77,178 | 0.0026% | 0 | 56,170 |
| 15591568 | rs200973492 | A | G | p.Val488Ala | c.1463T>C | 22.6 | 1 | 158,104 | 6.32E-06 | 1 | 11,663 | 0.0086% | 0 | 49,072 |
| 15582298 | rs41303171 | T | C | p.Asn720Asp | c.2158A>G | 15.09 | 2732 | 166,545 | 1.64E-02 | | | | 923 | 54,257 |
| Subtotal (p.Asn720Asp - not at SARS-CoV-2 Spike Protein Binding Interface) | | | | | | | 2732 | | | | | | | |
| Subpopulation-specific Prevalence (at least one p.Asn720Asp allele) | | | | | | | | | | | | | | |
| **Subtotal (Rare Missense SNVs at SARS-CoV-2 Spike Protein Binding Interface)** | | | | | | | 759 | | | | | | | |
| **Subpopulation-specific Prevalence (at least one rare missense SNVs at SARS-CoV-2 Spike Protein binding interface)** | | | | | | | | | 0.004142537 | | | | | |

| Minor Allele Frequency (MAF) (All Males in gnomAD Database) | Alternate Allele Count (All Females in gnomAD Database) | Total Allele Number (All Females in gnomAD Database) | Minor Allele Frequency (MAF) (All Females in gnomAD Database) | Alternate Allele Count Ashkenazi Jewish Ancestry | Total Allele Number Ashkenazi Jewish Ancestry | Minor Allele Frequency (MAF) Ashkenazi Jewish Ancestry | Alternate Allele Count Ashkenazi Jewish Males | Total Allele Number Ashkenazi Jewish Males | Minor Allele Frequency (MAF) Ashkenazi Jewish Males | Alternate Allele Count Ashkenazi Jewish Females | Total Allele Number Ashkenazi Jewish Females | Minor Allele Frequency (MAF) Ashkenazi Jewish Females | Alternate Allele Count North-Western European Ancestry | Total Allele Number North-Western European Ancestry |
|---|---|---|---|---|---|---|---|---|---|---|---|---|---|---|
| 1.48E-05 | 0 | 115,566 | 0.00E+00 | 0 | 7,485 | 0 | 0 | 2,585 | 0 | 0 | 4,900 | 0 | 0 | 30,821 |
| 3.82E-03 | 469 | 115,564 | 4.06E-03 | 90 | 7,483 | 0.012027262 | 37 | 2,583 | 0.014324429 | 53 | 4,900 | 0.010816327 | 208 | 30,823 |
| 0.00E+00 | 2 | 115,566 | 1.73E-05 | 0 | 7,485 | 0 | 0 | 2,585 | 0 | 0 | 4,900 | 0 | 0 | 30,823 |
| 1.47E-05 | 2 | 115,564 | 1.73E-05 | 0 | 7,482 | 0 | 0 | 2,582 | 0 | 0 | 4,900 | 0 | 1 | 30,826 |
| 4.43E-05 | 3 | 115,562 | 2.60E-05 | 0 | 7,479 | 0 | 0 | 2,579 | 0 | 0 | 4,900 | 0 | 0 | 30,818 |
| 0.00E+00 | 3 | 115,562 | 2.60E-05 | 0 | 7,480 | 0 | 0 | 2,580 | 0 | 0 | 4,900 | 0 | 0 | 30,800 |
| 0.00E+00 | 1 | 115,562 | 8.65E-06 | 0 | 7,477 | 0 | 0 | 2,577 | 0 | 0 | 4,900 | 0 | 0 | 30,763 |
| 0.00E+00 | 2 | 115,522 | 1.73E-05 | 0 | 7,471 | 0 | 0 | 2,571 | 0 | 0 | 4,900 | 0 | 0 | 30,727 |
| 0.00E+00 | 1 | 115,522 | 8.66E-06 | 0 | 7,470 | 0 | 0 | 2,570 | 0 | 0 | 4,900 | 0 | 0 | 30,731 |
| 2.98E-05 | 0 | 115,534 | 0.00E+00 | 0 | 7,477 | 0 | 0 | 2,577 | 0 | 0 | 4,900 | 0 | 0 | 30,731 |
| 0.00E+00 | 1 | 115,394 | 8.67E-06 | 0 | 7,441 | 0 | 0 | 2,543 | 0 | 0 | 4,898 | 0 | 0 | 30,397 |
| 0.00E+00 | 5 | 115,420 | 4.33E-05 | 0 | 7,448 | 0 | 0 | 2,550 | 0 | 0 | 4,898 | 0 | 1 | 30,440 |
| 0.00E+00 | 1 | 114,618 | 8.72E-06 | 0 | 7,025 | 0 | 0 | 2,159 | 0 | 0 | 4,866 | 0 | 0 | 28,562 |
| 0.00E+00 | 2 | 114,140 | 1.75E-05 | 0 | 6,759 | 0 | 0 | 1,929 | 0 | 0 | 4,830 | 0 | 1 | 27,814 |
| 0.00E+00 | 1 | 109,032 | 9.17E-06 | 0 | 6,523 | 0 | 0 | 1,785 | 0 | 0 | 4,738 | 0 | 0 | 24,554 |
| 1.70E-02 | 1809 | 112,288 | 1.61E-02 | 106 | 6,316 | 0.016782774 | 28 | 1,650 | 0.016969697 | 78 | 4,666 | 0.016716674 | 746 | 27,700 |
| 1.70% | | | 1.61% | | | | | | 1.70% | | | 1.67% | | |
| 1.70% | | | 3.22% | | | | | | 1.70% | | | 3.34% | | |
| 0.39% | | | 0.43% | | | | | | 1.43% | | | 1.08% | | |
| 0.39% | | | 0.85% | | | | | | 1.43% | | | 2.16% | | |

| Minor Allele Frequency (MAF) North-Western European Ancestry | Alternate Allele Count Non-Finnish Europeans | Total Allele Number Non-Finnish Europeans | Minor Allele Frequency (MAF) Non-Finnish Europeans | Alternate Allele Count Non-Finnish European Males | Total Allele Number Non-Finnish European Males | Minor Allele Frequency (MAF) Non-Finnish European Males | Alternate Allele Count Non-Finnish European Females | Total Allele Number Non-Finnish European Females | Minor Allele Frequency (MAF) Non-Finnish European Females | Alternate Allele Count Non-Finnish Europeans (of Otherwise indeterminate Ancestry) | Total Allele Number Non-Finnish Europeans (of Otherwise indeterminate Ancestry) | Minor Allele Frequency (MAF) Non-Finnish Europeans (of Otherwise indeterminate Ancestry) | Alternate Allele Count Swedish Ancestry | Total Allele Number Swedish Ancestry |
|---|---|---|---|---|---|---|---|---|---|---|---|---|---|---|
| 0 | 1 | 81,870 | 1.22145E-05 | 1 | 31,716 | 3.15298E-05 | 0 | 50,154 | 0 | 0 | 22,665 | 0 | 1 | 19,164 |
| 0.006748208 | 483 | 81,876 | 0.005899165 | 180 | 31,724 | 0.005673938 | 303 | 50,152 | 0.006041633 | 141 | 22,669 | 0.006219948 | 99 | 19,164 |
| 0 | 0 | 81,875 | 0 | 0 | 31,721 | 0 | 0 | 50,154 | 0 | 0 | 22,668 | 0 | 0 | 19,164 |
| 3.24401E-05 | 1 | 81,891 | 1.22114E-05 | 0 | 31,739 | 0 | 1 | 50,152 | 1.99394E-05 | 0 | 22,682 | 0 | 0 | 19,163 |
| 0 | 0 | 81,876 | 0 | 0 | 31,726 | 0 | 0 | 50,150 | 0 | 0 | 22,675 | 0 | 0 | 19,162 |
| 0 | 0 | 81,854 | 0 | 0 | 31,700 | 0 | 0 | 50,154 | 0 | 0 | 22,669 | 0 | 0 | 19,163 |
| 0 | 0 | 81,803 | 0 | 0 | 31,649 | 0 | 0 | 50,154 | 0 | 0 | 22,654 | 0 | 0 | 19,164 |
| 0 | 0 | 81,741 | 0 | 0 | 31,605 | 0 | 0 | 50,136 | 0 | 0 | 22,641 | 0 | 0 | 19,150 |
| 0 | 0 | 81,743 | 0 | 0 | 31,605 | 0 | 0 | 50,138 | 0 | 0 | 22,636 | 0 | 0 | 19,154 |
| 0 | 0 | 81,689 | 0 | 2 | 31,551 | 0 | 0 | 50,138 | 0 | 0 | 22,626 | 0 | 0 | 19,144 |
| 0 | 0 | 81,315 | 0 | 0 | 31,231 | 0 | 0 | 50,084 | 0 | 0 | 22,547 | 0 | 0 | 19,160 |
| 3.28515E-05 | 4 | 81,369 | 4.91588E-05 | 0 | 31,277 | 0 | 4 | 50,092 | 7.98531E-05 | 2 | 22,556 | 8.86682E-05 | 0 | 19,161 |
| 0 | 1 | 78,468 | 1.2744E-05 | 0 | 28,644 | 0 | 1 | 49,824 | 2.00706E-05 | 1 | 21,742 | 4.59939E-05 | 0 | 19,077 |
| 3.59531E-05 | 2 | 77,178 | 2.59141E-05 | 0 | 27,538 | 0 | 2 | 49,640 | 4.02901E-05 | 1 | 21,311 | 4.69241E-05 | 0 | 19,036 |
| 0 | 0 | 73,143 | 0 | 0 | 25,105 | 0 | 0 | 48,038 | 0 | 0 | 20,626 | 0 | 0 | 18,951 |
| 0.026931408 | 1971 | 76,316 | 0.025826825 | 678 | 27,288 | 0.024846086 | 1293 | 49,028 | 0.026372685 | 561 | 20,875 | 0.026874251 | 542 | 18,843 |
| | | | | | | 2.48% | | | 2.64% | | | | | |
| | | | | | | 2.48% | | | 5.27% | | | | | |
| | | | | | | 0.58% | | | 0.62% | | | | | |
| | | | | | | 0.58% | | | 1.24% | | | | | |

| Minor Allele Frequency (MAF) Swedish Ancestry | Alternate Allele Count Estonian Ancestry | Total Allele Number Estonian Ancestry | Minor Allele Frequency (MAF) Estonian Ancestry | Alternate Allele Count Bulgarian Ancestry | Total Allele Number Bulgarian Ancestry | Minor Allele Frequency (MAF) Bulgarian Ancestry | Alternate Allele Count Southern Europeans | Total Allele Number Southern Europeans | Minor Allele Frequency (MAF) Southern Europeans | Alternate Allele Count Samples of Uncertain Ancestry | Total Allele Number Samples of Uncertain Ancestry | Minor Allele Frequency (MAF) Samples of Uncertain Ancestry | Alternate Allele Count Males of Uncertain Ancestry |
|---|---|---|---|---|---|---|---|---|---|---|---|---|---|
| 5.21812E-05 | 0 | 197 | 0 | 0 | 1,983 | 0 | 0 | 7,040 | 0 | 0 | 4,527 | 0 | 0 |
| 0.005165936 | 1 | 197 | 0.005076142 | 10 | 1,983 | 0.005042864 | 24 | 7,040 | 0.003409091 | 17 | 4,528 | 0.003754417 | 5 |
| 0 | 0 | 197 | 0 | 0 | 1,983 | 0 | 0 | 7,040 | 0 | 0 | 4,529 | 0 | 0 |
| 0 | 0 | 197 | 0 | 0 | 1,983 | 0 | 0 | 7,040 | 0 | 0 | 4,529 | 0 | 0 |
| 0 | 0 | 197 | 0 | 0 | 1,983 | 0 | 0 | 7,041 | 0 | 0 | 4,527 | 0 | 0 |
| 0 | 0 | 197 | 0 | 0 | 1,982 | 0 | 0 | 7,043 | 0 | 0 | 4,523 | 0 | 0 |
| 0 | 0 | 197 | 0 | 0 | 1,983 | 0 | 0 | 7,042 | 0 | 0 | 4,516 | 0 | 0 |
| 0 | 0 | 197 | 0 | 0 | 1,982 | 0 | 0 | 7,044 | 0 | 0 | 4,524 | 0 | 0 |
| 0 | 0 | 197 | 0 | 0 | 1,981 | 0 | 0 | 7,044 | 0 | 0 | 4,524 | 0 | 0 |
| 0 | 0 | 197 | 0 | 0 | 1,981 | 0 | 2 | 7,040 | 0.000284091 | 0 | 4,514 | 0 | 0 |
| 0 | 0 | 195 | 0 | 0 | 1,982 | 0 | 0 | 7,034 | 0 | 0 | 4,443 | 0 | 0 |
| 0 | 0 | 196 | 0 | 0 | 1,982 | 0 | 1 | 7,034 | 0.000142167 | 1 | 4,441 | 0.000225175 | 0 |
| 0 | 0 | 189 | 0 | 0 | 1,968 | 0 | 0 | 6,930 | 0 | 0 | 4,222 | 0 | 0 |
| 0 | 0 | 186 | 0 | 0 | 1,956 | 0 | 0 | 6,875 | 0 | 0 | 4,129 | 0 | 0 |
| 0 | 0 | 178 | 0 | 0 | 1,953 | 0 | 0 | 6,881 | 0 | 0 | 3,680 | 0 | 0 |
| 0.028763997 | 7 | 180 | 0.038888889 | 31 | 1,936 | 0.016012397 | 84 | 6,782 | 0.012385727 | 50 | 4,016 | 0.012450199 | 20 |

| Total Allele Number Males of Uncertain Ancestry | Minor Allele Frequency (MAF) Males of Uncertain Ancestry | Alternate Allele Count Females of Uncertain Ancestry | Total Allele Number Females of Uncertain Ancestry | Minor Allele Frequency (MAF) Females of Uncertain Ancestry | Alternate Allele Count Latino Ancestry | Total Allele Number Latino Ancestry | Minor Allele Frequency (MAF) Latino Ancestry | Alternate Allele Count Males of Latino Ancestry | Total Allele Number Males of Latino Ancestry | Minor Allele Frequency (MAF) Males of Latino Ancestry | Alternate Allele Count Females of Latino Ancestry | Total Allele Number Females of Latino Ancestry | Minor Allele Frequency (MAF) Females of Latino Ancestry | Alternate Allele Count Africans |
|---|---|---|---|---|---|---|---|---|---|---|---|---|---|---|
| 1,599 | 0 | 0 | 2,928 | 0 | 0 | 27,386 | 0 | 0 | 7,118 | 0 | 0 | 20,268 | 0 | 0 |
| 1,600 | 0.003125 | 12 | 2,928 | 0.004098361 | 90 | 27,389 | 0.00328599 | 20 | 7,121 | 0.00280859 | 70 | 20,268 | 0.00345372 | 13 |
| 1,601 | 0 | 0 | 2,928 | 0 | 2 | 27,385 | 7.30327E-05 | 0 | 7,117 | 0 | 2 | 20,268 | 9.86777E-05 | 0 |
| 1,601 | 0 | 0 | 2,928 | 0 | 0 | 27,394 | 0 | 0 | 7,126 | 0 | 0 | 20,268 | 0 | 0 |
| 1,599 | 0 | 0 | 2,928 | 0 | 0 | 27,387 | 0 | 0 | 7,119 | 0 | 0 | 20,268 | 0 | 1 |
| 1,595 | 0 | 0 | 2,928 | 0 | 3 | 27,349 | 0.000109693 | 0 | 7,083 | 0 | 3 | 20,266 | 0.000148031 | 0 |
| 1,588 | 0 | 0 | 2,928 | 0 | 0 | 27,320 | 0 | 0 | 7,056 | 0 | 0 | 20,264 | 0 | 0 |
| 1,598 | 0 | 0 | 2,926 | 0 | 0 | 27,302 | 0 | 0 | 7,042 | 0 | 0 | 20,260 | 0 | 2 |
| 1,598 | 0 | 0 | 2,926 | 0 | 0 | 27,310 | 0 | 0 | 7,052 | 0 | 0 | 20,258 | 0 | 0 |
| 1,588 | 0 | 0 | 2,926 | 0 | 0 | 27,267 | 0 | 0 | 7,007 | 0 | 0 | 20,260 | 0 | 0 |
| 1,519 | 0 | 0 | 2,924 | 0 | 0 | 26,817 | 0 | 0 | 6,615 | 0 | 0 | 20,202 | 0 | 1 |
| 1,519 | 0 | 1 | 2,922 | 0.000342231 | 0 | 26,833 | 0 | 0 | 6,625 | 0 | 0 | 20,208 | 0 | 0 |
| 1,332 | 0 | 0 | 2,890 | 0 | 0 | 25,656 | 0 | 0 | 5,720 | 0 | 0 | 19,936 | 0 | 0 |
| 1,247 | 0 | 0 | 2,882 | 0 | 0 | 25,097 | 0 | 0 | 5,313 | 0 | 0 | 19,784 | 0 | 0 |
| 998 | 0 | 0 | 2,682 | 0 | 0 | 22,256 | 0 | 0 | 3,870 | 0 | 0 | 18,386 | 0 | 1 |
| 1,210 | 0.016528926 | 30 | 2,806 | 0.010691376 | 64 | 23,747 | 0.00269508 | 12 | 4,601 | 0.00260813 | 52 | 19,146 | 0.00271597 | 39 |
| | 1.65% | | | 1.07% | | | | | | 0.26% | | | 0.27% | |
| | 1.65% | | | 2.14% | | | | | | 0.26% | | | 0.54% | |
| | 0.31% | | | 0.44% | | | | | | 0.28% | | | 0.37% | |
| | 0.31% | | | 0.89% | | | | | | 0.28% | | | 0.74% | |

| Total Allele Number Africans | Minor Allele Frequency (MAF) Africans | Alternate Allele Count African Males | Total Allele Number African Males | Minor Allele Frequency (MAF) African Males | Alternate Allele Count African Females | Total Allele Number African Females | Minor Allele Frequency (MAF) African Females | Alternate Allele Count South Asians | Total Allele Number South Asians | Minor Allele Frequency (MAF) South Asians | Alternate Allele Count South Asian Males | Total Allele Number South Asian Males | Minor Allele Frequency (MAF) South Asian Males | Alternate Allele Count South Asian Females |
|---|---|---|---|---|---|---|---|---|---|---|---|---|---|---|
| 13,161 | 0 | 0 | 3,091 | 0 | 0 | 10,070 | 0 | 0 | 19,045 | 0 | 0 | 11,501 | 0 | 0 |
| 13,162 | 0.000987692 | 4 | 3,092 | 0.001293661 | 9 | 10,070 | 0.000893744 | 25 | 19,044 | 0.001312749 | 9 | 11,500 | 0.000782609 | 16 |
| 13,162 | 0 | 0 | 3,092 | 0 | 0 | 10,070 | 0 | 0 | 19,034 | 0 | 0 | 11,490 | 0 | 0 |
| 13,162 | 0 | 0 | 3,092 | 0 | 0 | 10,070 | 0 | 0 | 19,061 | 0 | 0 | 11,517 | 0 | 0 |
| 13,163 | 7.59705E-05 | 0 | 3,093 | 0 | 1 | 10,070 | 9.93049E-05 | 0 | 19,055 | 0 | 0 | 11,511 | 0 | 0 |
| 13,160 | 0 | 0 | 3,090 | 0 | 0 | 10,070 | 0 | 0 | 19,030 | 0 | 0 | 11,486 | 0 | 0 |
| 13,157 | 0 | 0 | 3,087 | 0 | 0 | 10,070 | 0 | 1 | 18,988 | 5.26648E-05 | 0 | 11,444 | 0 | 1 |
| 13,148 | 0.000152114 | 0 | 3,078 | 0 | 2 | 10,070 | 0.00019861 | 0 | 18,809 | 0 | 0 | 11,267 | 0 | 0 |
| 13,143 | 0 | 0 | 3,073 | 0 | 0 | 10,070 | 0 | 1 | 18,823 | 5.31265E-05 | 0 | 11,283 | 0 | 1 |
| 13,147 | 0 | 0 | 3,077 | 0 | 0 | 10,070 | 0 | 0 | 16,692 | 0 | 0 | 11,152 | 0 | 0 |
| 13,058 | 7.65814E-05 | 0 | 2,994 | 0 | 1 | 10,064 | 9.93641E-05 | 0 | 18,541 | 0 | 0 | 11,021 | 0 | 0 |
| 13,070 | 0 | 0 | 3,002 | 0 | 0 | 10,068 | 0 | 0 | 18,577 | 0 | 0 | 11,047 | 0 | 0 |
| 12,788 | 0 | 0 | 2,736 | 0 | 0 | 10,052 | 0 | 0 | 16,880 | 0 | 0 | 9,450 | 0 | 0 |
| 12,716 | 0 | 0 | 2,662 | 0 | 0 | 10,054 | 0 | 0 | 15,981 | 0 | 0 | 8,585 | 0 | 0 |
| 11,663 | 8.57412E-05 | 0 | 2,115 | 0 | 1 | 9,548 | 0.000104734 | 0 | 14,330 | 0 | 0 | 7,464 | 0 | 0 |
| 12,775 | 0.003052838 | 10 | 2,795 | 0.003577818 | 29 | 9,980 | 0.002905812 | 66 | 15,207 | 0.004340107 | 40 | 7,937 | 0.005039688 | 26 |
| | | | | 0.36% | | | 0.29% | | | | | | 0.50% | |
| | | | | 0.36% | | | 0.58% | | | | | | 0.50% | |
| | | | | 0.13% | | | 0.14% | | | | | | 0.08% | |
| | | | | 0.13% | | | 0.28% | | | | | | 0.08% | |

| Total Allele Number South Asian Females | Minor Allele Frequency (MAF) South Asian Females | Alternate Allele Count Finnish | Total Allele Number Finnish | Minor Allele Frequency (MAF) Finnish | Alternate Allele Count Finnish Males | Total Allele Number Finnish Males | Minor Allele Frequency (MAF) Finnish Males | Alternate Allele Count Finnish Females | Total Allele Number Finnish Females | Minor Allele Frequency (MAF) Finnish Females | Alternate Allele Count East Asians | Total Allele Number East Asians | Minor Allele Frequency (MAF) East Asians |
|---|---|---|---|---|---|---|---|---|---|---|---|---|---|
| 7,544 | 0 | 0 | 16,008 | 0 | 0 | 5,634 | 0 | 0 | 10,374 | 0 | 0 | 13,844 | 0 |
| 7,544 | 0.002120891 | 9 | 16,008 | 0.000562219 | 4 | 5,634 | 0.000709975 | 5 | 10,374 | 0.000481974 | 1 | 13,840 | 7.22543E-05 |
| 7,544 | 0 | 0 | 16,006 | 0 | 0 | 5,632 | 0 | 0 | 10,374 | 0 | 0 | 13,843 | 0 |
| 7,544 | 0 | 0 | 16,008 | 0 | 0 | 5,634 | 0 | 0 | 10,374 | 0 | 2 | 13,847 | 0.000144436 |
| 7,544 | 0 | 5 | 16,006 | 0.000312383 | 3 | 5,632 | 0.00053267 | 2 | 10,374 | 0.00019279 | 0 | 13,846 | 0 |
| 7,544 | 0 | 0 | 16,003 | 0 | 0 | 5,631 | 0 | 0 | 10,372 | 0 | 0 | 13,834 | 0 |
| 7,544 | 0.000132556 | 0 | 16,003 | 0 | 0 | 5,629 | 0 | 0 | 10,374 | 0 | 0 | 13,824 | 0 |
| 7,542 | 0 | 0 | 15,942 | 0 | 0 | 5,580 | 0 | 0 | 10,362 | 0 | 0 | 13,821 | 0 |
| 7,540 | 0.000132626 | 0 | 15,953 | 0 | 0 | 5,589 | 0 | 0 | 10,364 | 0 | 0 | 13,826 | 0 |
| 7,540 | 0 | 0 | 15,953 | 0 | 0 | 5,581 | 0 | 0 | 10,372 | 0 | 0 | 13814 | 0 |
| 7,520 | 0 | 0 | 15,989 | 0 | 0 | 5,615 | 0 | 0 | 10,374 | 0 | 0 | 13,662 | 0 |
| 7,530 | 0 | 0 | 15,990 | 0 | 0 | 5,616 | 0 | 0 | 10,374 | 0 | 0 | 13,673 | 0 |
| 7,430 | 0 | 0 | 15,739 | 0 | 0 | 5,411 | 0 | 0 | 10,328 | 0 | 0 | 13,069 | 0 |
| 7,396 | 0 | 0 | 15,563 | 0 | 0 | 5,283 | 0 | 0 | 10,280 | 0 | 0 | 12,887 | 0 |
| 6,866 | 0 | 0 | 15,498 | 0 | 0 | 5,224 | 0 | 0 | 10,274 | 0 | 0 | 11,011 | 0 |
| 7,270 | 0.003576341 | 436 | 15,384 | 0.028341134 | 135 | 5,132 | 0.026305534 | 301 | 10,252 | 0.029360125 | 0 | 12,784 | 0 |
| | 0.36% | | | | | | 2.63% | | | 2.94% | | | |
| | 0.72% | | | | | | 2.63% | | | 5.87% | | | |
| | 0.24% | | | | | | 0.12% | | | 0.07% | | | |
| | 0.48% | | | | | | 0.12% | | | 0.13% | | | |

| Alternate Allele Count East Asian Males | Total Allele Number East Asian Males | Minor Allele Frequency (MAF) East Asian Males | Alternate Allele Count East Asian Females | Total Allele Number East Asian Females | Minor Allele Frequency (MAF) East Asian Females | Alternate Allele Count Japanese Ancestry | Total Allele Count Japanese Ancestry | Minor Allele Frequency (MAF) Japanese Ancestry | Alternate Allele Count Korean Ancestry | Total Allele Count Korean Ancestry | Minor Allele Frequency (MAF) Korean Ancestry | Alternate Allele Count East Asian Ancestry (non-Korean, non-Japanese) | Total Allele Number East Asian Ancestry (non-Korean, non-Japanese) | Minor Allele Frequency (MAF) East Asian Ancestry (non-Korean, non-Japanese) |
|---|---|---|---|---|---|---|---|---|---|---|---|---|---|---|
| 0 | 4,516 | 0 | 0 | 9,328 | 0 | 0 | 122 | 0 | 0 | 2,887 | 0 | 0 | 10,835 | 0 |
| 0 | 4,512 | 0 | 1 | 9,328 | 0.000107204 | 0 | 122 | 0 | 0 | 2,887 | 0 | 1 | 10,831 | 9.23276E-05 |
| 0 | 4,515 | 0 | 0 | 9,328 | 0 | 0 | 122 | 0 | 0 | 2,887 | 0 | 0 | 10,834 | 0 |
| 1 | 4,519 | 0.000221288 | 1 | 9,328 | 0.000107204 | 0 | 122 | 0 | 0 | 2,887 | 0 | 2 | 10,838 | 0.000184536 |
| 0 | 4,518 | 0 | 0 | 9,328 | 0 | 0 | 122 | 0 | 0 | 2,887 | 0 | 0 | 10,837 | 0 |
| 0 | 4,506 | 0 | 0 | 9,328 | 0 | 0 | 122 | 0 | 0 | 2,887 | 0 | 0 | 10,825 | 0 |
| 0 | 4,496 | 0 | 0 | 9,328 | 0 | 0 | 122 | 0 | 0 | 2,885 | 0 | 0 | 10,817 | 0 |
| 0 | 4,495 | 0 | 0 | 9,326 | 0 | 0 | 122 | 0 | 0 | 2,884 | 0 | 0 | 10,815 | 0 |
| 0 | 4,500 | 0 | 0 | 9,326 | 0 | 0 | 122 | 0 | 0 | 2,883 | 0 | 0 | 10,821 | 0 |
| 0 | 4486 | 0 | 0 | 9328 | 0 | 0 | 122 | 0 | 0 | 2,885 | 0 | 0 | 10,807 | 0 |
| 0 | 4,334 | 0 | 0 | 9,328 | 0 | 0 | 122 | 0 | 0 | 2,817 | 0 | 0 | 10,723 | 0 |
| 0 | 4,345 | 0 | 0 | 9,328 | 0 | 0 | 122 | 0 | 0 | 2,821 | 0 | 0 | 10,730 | 0 |
| 0 | 3,777 | 0 | 0 | 9,292 | 0 | 0 | 120 | 0 | 0 | 2,695 | 0 | 0 | 10,254 | 0 |
| 0 | 3,613 | 0 | 0 | 9,274 | 0 | 0 | 120 | 0 | 0 | 2,662 | 0 | 0 | 10,105 | 0 |
| 0 | 2,511 | 0 | 0 | 8,500 | 0 | 0 | 120 | 0 | 0 | 2,232 | 0 | 0 | 8,659 | 0 |
| 0 | 3,644 | 0 | 0 | 9,140 | 0 | 0 | 119 | 0 | 0 | 2,798 | 0 | 0 | 9,867 | 0 |
| | | 0.00% | | | 0.00% | | | | | | | | | |
| | | 0.00% | | | 0.00% | | | | | | | | | |
| | | 0.02% | | | 0.02% | | | | | | | | | |
| | | 0.02% | | | 0.04% | | | | | | | | | |
