## Supplementary Table 2 for "ACE 2 Coding Variants: A Potential X-linked Risk Factor for COVID-19 Disease"

**Variants Ordered by Position**

| resWild | residue# | resMut | ddGbind |
|---|---|---|---|
| K | 26 | E | 0.721204 |
| K | 26 | R | -1.12335 |
| T | 27 | A | 0.907301 |
| E | 35 | K | 0.0219484 |
| E | 37 | K | 1.23186 |
| F | 40 | L | 0.0376725 |
| S | 43 | R | -0.536781 |
| M | 82 | I | -0.627869 |
| P | 84 | T | 0.00646006 |
| G | 326 | E | -0.588113 |
| E | 329 | G | 0.78436 |
| G | 352 | V | 0.424471 |
| D | 355 | N | 0.0205439 |
| V | 488 | A | 0.286061 |
| N | 720 | D | -0.119618 |

**Variants Ordered by Rank**

| resWild | residue# | resMut | ddGbind |
|---|---|---|---|
| E | 37 | K | 1.23186 |
| T | 27 | A | 0.907301 |
| E | 329 | G | 0.78436 |
| K | 26 | E | 0.721204 |
| G | 352 | V | 0.424471 |
| V | 488 | A | 0.286061 |
| F | 40 | L | 0.0376725 |
| E | 35 | K | 0.0219484 |
| D | 355 | N | 0.0205439 |
| P | 84 | T | 0.00646006 |
| N | 720 | D | -0.119618 |
| S | 43 | R | -0.536781 |
| G | 326 | E | -0.588113 |
| M | 82 | I | -0.627869 |
| K | 26 | R | -1.12335 |
